# An LPS-dephosphorylating alkaline phosphatase of PhoA family from the marine bacterium *Cobetia amphilecti* KMM 296 and multiplicity of alkaline phosphatase families in *Cobetia* spp

**DOI:** 10.1101/2024.01.29.577745

**Authors:** Larissa Balabanova, Svetlana Bakholdina, Nina Buinovskaya, Yulia Noskova, Oksana Kolpakova, Vanessa Vlasova, Georgii Bondarev, Aleksandra Seitkalieva, Oksana Son, Liudmila Tekutyeva

**Affiliations:** Laboratory of Marine Biochemistry, G.B. Elyakov Pacific Institute of Bioorganic Chemistry, Far Eastern Branch, Russian Academy of Sciences, Prospect 100-Letya Vladivostoka 152, 690022 Vladivostok, Russia; Advanced Engineering School, Institute of Biotechnology, Bioengineering and Food Systems, Far Eastern Federal University, 10 Ajax Bay, Russky Island, 690922 Vladivostok, Russia; Medical and Genetic Centre, Central office, Laboratory of MGC “Genomed”, 8 Korolenko str., 107014 Moscow, Russia

**Keywords:** *Cobetia* spp. genomes, alkaline phosphatase PhoA, PhoD, PhoX, PafA family, endotoxin, LPS phosphorus hydrolysis, organophosphate degradation, sulfated extracellular polysaccharides, ectoine biosynthetic gene cluster

## Abstract

A highly active alkaline phosphatase (ALP) from the mollusk strain of the marine bacterium *Cobetia amphilecti* KMM 296 (CmAP) was found to remove phosphorus from the *Escherichia coli* lipopolysaccharides (LPS). Phylogenetic analysis of the amino acid sequences of ALPs found in 36 available *Cobetia* genomes revealed that CmAP and its homologues from nine strains clustered together with the human and squid LPS-detoxifying enzymes. Each strain of the genus *Cobetia* has a variety of ALPs mostly of the PhoD and PhoX families. The PhoA gene encoding for the CmAP-like ALP is characteristic for the subspecies *C. amphilecti*, with a complete set of four ALP families, including PafA and two PhoD structures (5 genes). However, a single strain of the species *Cobetia crustatorum* JO1^T^ from fermented shrimp, phylogenetically distant from *C. amphilecti* and *C. marina*, among four ALPs contains a CmAP homologue carrying an inactive mutation. Apparently, the multiplicity of ALPs in bacteria of the genus *Cobetia* is a trait of incredible adaptation to a phosphorus-depleted environment and a specialty of organophosphate destructor in eco-niches to which they once emerged, including *Zostera* spp. roots. The ALP clusterization and an identity level of the genus-specific biosynthetic genes encoding for ectoine and polyketide cluster T1PKS, responsible for sulfated extracellular polysaccharide synthesis, coincide with a new whole genome-based taxonomic classification of the genus *Cobetia*. The LPS-dephosphorylating property of the PhoA family *C. amphilecti* ALP CmAP may be used in the development of anti-inflammatory drugs.

## 1. Introduction

Alkaline phosphatases (ALPs) (EC. 3.1.3.1) belong to nonspecific metal-dependent ectoenzymes that catalyze the hydrolysis and transphosphorylation of complex phosphoric acid monoesters in the environment by a mechanism involving the formation of a covalent phosphoserine intermediate and the release of inorganic phosphate (P_i_) and alcohol (or phenol) at alkaline pH values [1-3]. Alkaline phosphatases are widespread in nature from bacteria to humans. In mammals, ALPs are represented as a group of isoenzymes expressed in different tissues and differ in physicochemical properties and physiological functions that have not yet been fully understood [1,4,5]. The most important function of the intestinal ALP isoenzyme (IAP) is dephosphorylation of Gram-negative bacterial endotoxins (lipopolysaccharides, LPS) and other inflammatory mediators responsible for chronic systemic diseases, as well as the maintenance of intestinal microbial homeostasis and intestinal barrier function. Thus, deficiency of IAP is responsible for a number of pathologies, including metabolic syndrome, liver fibrosis, coronary heart disease, osteoporosis, and aging in general [4,6-8]. The increased levels of non-dephosphorylated LPS in humans correlate with an increased risk of liver and colorectal cancer [9].

The tissue-specific ALP of the columnar epithelium in animals and invertebrates has also become associated with an interaction of the host and its microbiome due to its ability to dephosphorylate lipid A, a component of LPS [10]. Moreover, the recognition of bacterial LPS and their dephosphorylation are rather ancient functions of this family of enzymes and belong to the innate immune system of multicellular organisms, which approved in experiments on colonization of the light organ (photophore) of the squid *Euprymna scolopes* by its symbiotic luminescent bacterium *Vibrio fischeri* [11]. An increase in the level of dephosphorylated LPS by sunset is a signal for the bacteria to increase the population size and, accordingly, the intensity of photophore luminescence. At the same time, the LPS dephosphorylation protects the invertebrate from excessive inflammation and destruction of its own tissues under the influence of endotoxin, the phosphorylated lipid A [11].

In bacteria, ALPs play a major role in utilizing organic phosphates as an alternative source of the vital macronutrient, phosphorus (P), under deficiency of P_i_ in the environment [12-14]. The bacterial ALPs catalyze the hydrolysis of sugar phosphates, DNA and RNA (5’-, 3’-ends), nucleotide mono-, di-, and triphosphates (dNMP, dNDP, dNTP), lipid phosphatidates, polyphosphates (polyP), and pyrophosphates, which are prevalent in the environment enriched in organics [3,15-17]. However, some APSs were found to catalyze the cleavage of sulfate and phosphate di- and triesthers, including neurotoxins with P-O-C bonds, due to their evolutionarily emerged catalytic promiscuity, thus possessing an ecological relevance [13,18,19]. In addition, ALPs nonspecifically dephosphorylate some proteins, many of which are the part of cell signaling transduction [20,21]. Finally, microbial ALPs are of global biogeochemical importance, being among the most abundant enzymes in soils and the World’s Ocean [22-26]. Alkaline phosphatases have been shown to be involved in the bacterial competing with fungi by regulating their metabolic pathways [27]; to induce the bacterial biofilm and invertebrate exoskeleton’s growth and mineralization [15,28,29]; to participate in remediation of heavy metals and organic pollutants [13,14,30]. Alkaline phosphatases are widely distributed among marine bacteria that extract P_i_ from dissolved organic P-containing compounds in the global ocean [12-14,22,25,31]. In addition, marine bacteria and diatom algae can accumulate P_i_ and polyP, with their subsequent release from the cells into the environment and a concomitant increase in the level of phosphatase activity [12,32]. Thus, microorganisms take an active part in the enzymatic induction of nucleation and crystal growth of such minerals as apatite and phosphorites [24-26].

Currently, four large structural families of prokaryotic ALPs are known, namely: PhoA, PhoD, PhoX, and PafA [14,22,23,26]. They differ from each other in structure, mechanism of enzymatic action, subcellular localization, substrate specificity and dependence of the manifestation of their activity on various metal ions, temperature and pH ranges [14,31,33]. Multicellular organisms produce the Mg^2+^/Zn^2+^-dependent ALPs structurally belonging to the PhoA family, which includes the ALPs of the model microorganism *E. coli* and mammals, therefore they are better characterized and considered as the classical ALPs [1-2,16,29]. Despite conservative the main characteristics of the catalytic mechanism, the mammalian ALPs have higher specific activity and effectiveness, lower values of the Michaelis constant (*K*_M_), as well as a more alkaline pH optimum compared to bacterial ALPs [1]. However, the enzymes of some marine bacteria, for example, of *C. amphilecti* KMM 296 (CmAP), have the catalytic parameters comparable of the mammalian ALPs and significant enzymatic activity in the monomeric state that makes them attractive for use in biotechnology [33].

In this study, we used a quantitative phosphorus assay to determine a suggested enzymatic activity of the *C. amphilecti* KMM 296 alkaline phosphatase CmAP towards *E. coli* LPS. We identified the ALP structural family of the enzyme CmAP, as well as all homologous CmAP-like structures and non-homologous ALPs in all genomes of the *Cobetia* spp. strains available in the NCBI database. In order to perform phylogenetic analysis for the *Cobetia* spp. ALPs, the known LPS-dephosphorylating ALPs of human and invertebrate with an established function in modulating own microbiome were used. The species-specific identification of the *Cobetia* isolates was suggested based on the whole-genome ALP contents and biosynthetic gene profiles and identities in order to determine the difference between the PhoA-containing and non-containing bacteria. Analysis of the PhoA gene localization in the chromosomes was carried out to suggest its function and participation in metabolic pathways. The biochemical, ecological and therapeutic potential for the marine bacteria of the genus *Cobetia* were established.

## 2. Materials and Methods

### 2.1. Recombinant production of the C. amphilecti KMM 296 alkaline phosphatase CmAP

The recombinant protein CmAP was produced in the *E. coli* strain Rossetta DE3 using the recombinant plasmid Pho40 based on the pET-40b(+) vector (Novogen) including the full-length coding sequence of the mature alkaline phosphatase CmAP from the marine bacterium *C. amphilecti* KMM 296 as described earlier [33]. For heterologous expression, the competent cells of *E. coli* Rosetta (DE3) were transformed by the recombinant plasmid Pho40 and grown at 37 °C on the agar LB medium containing 25 μg/mL of kanamycin during the night. The recombinant clones were grown in 25 mL of the liquid LB medium containing 25 μg/mL of kanamycin at 200 rpm for 16 h at 28 °C. The cell cultures were placed in a fresh LB medium (1 L) containing kanamycin (25 μg/mL) and incubated at 37 °C on a shaker at 200 rpm, until the optical density at 600 nm was 0.6–0.8. After that, 0.2 mM isopropyl-β-D-thiogalactopyranoside (IPTG, Sigma-Aldrich, USA) was added to induce the recombinant gene expression, and incubation was continued at 16°C for 16-18 h at 200 rpm. Cells were pelleted via centrifugation at 4000×g rpm for 15 min at 8 °C, suspended in 250 mL of a buffer A containing 20 mM Tris-HCl (pH 9.0), 5 mM imidazole, 0.5 M NaCl, 20% glycerol (Sigma-Aldrich, USA) and subjected to an ultrasonic treatment by Bundeline SONOPULS HD 2070 (Berlin, Germany) to provide a complete release of the soluble recombinant protein from the *E. coli* periplasmic space.

After centrifugation of the disintegrated lysate at 11,000 rpm for 20 min, the supernatant was saturated with dry ammonium sulfate to 30%. The precipitate was removed by centrifugation (15 min at 11,000 rpm, 4 °C). The supernatant was collected and saturated with ammonium sulfate to 70% over an hour, resulting in precipitation of the recombinant protein. The precipitate was centrifuged for 15 min at 11,000 rpm at room temperature and resuspended in the buffer A. The resulting supernatant was applied to a metal affinity resin (Ni^2+^-IMAC-Sepharose, V column = 160 mL, GE Healthcare), equilibrated in the same buffer at the rate 1 mL/min. The column was washed with the buffer A, after which the recombinant protein was eluted with a linear gradient of 0.005– 0.5 M imidazole in the buffer A at an elution rate 2 mL/min. After that, dialysis was performed for 12 h against the buffer A containing 50% glycerol (Sigma-Aldrich, USA). The His-tag of the recombinant protein was removed during 12 h of incubation with 1-2 units of the enteropeptidase (L-HEP) per 1 mg of the recombinant protein with stirring. After rechromatography on the metal affinity sorbent, the unbound fractions were collected and applied to a Sours 15 Q ion exchange sorbent (column V = 8 mL, GE Healthcare) at a rate of 5 mL/min. Elution was carried out with a gradient of 0.15-1.0 M NaCl in the buffer A (without the salt) at the rate 2 mL/min. Fractions of the target protein were identified using the substrate *p*-nitrophenyl phosphate (*p*NPP) by the presence of ALP activity and by the molecular weight determined by SDS-PAGE electrophoresis [34].

### 2.2. Alkaline phosphatase activity assay

The standard assay for ALP activity was carried out in 500 μL of the reaction mixture containing 2 mM *p*NPP (Sigma-Aldrich, USA) in 0.1 M Tris–HCl buffer, pH 10.0, 0.2 M KCl at 37 °C for 30 min. The release of *p*-nitrophenol (ε = 18.5 mM/cm) was monitored at 405 nm after the addition of 2 mL 0.5 M NaOH (stop reagent). One unit of the ALP activity was defined as the quantity of the enzyme required to release 1.0 μM of *p*-nitrophenyl from *p*NPP in 1 min. The specific activity was calculated as units (U) per 1 mg of protein: U=(ΣV _reaction mixture + stop reagent_, mL × A_400_)/(18.3 _extinction coefficient_ × t _incubation_,min × V _protein solution_, mL × C _protein_, mg/mL).

### 2.3. Dephosphorylation activity assay towards E. coli LPS

The LPS from *E. coli* serotype 055:B5 (Sigma, USA) samples (1 and 0.2 mg/mL) were dissolved in 0.1 M Tris-HCl buffer, pH 7.7, containing 0.1 M KCl in three ways: 1) incubation at 24 °C for 12 h (sample N 1); 2) incubation at 37 °C for 12 h (sample N 2); 3) incubation at 24 °C for 12 h in the same buffer with the addition of organic solvent - triethylamine (TEA, 1μl/mL) to pH 10.0, a strong LPS dispersant (sample N 3).

Enzymatic activity of the recombinant alkaline phosphatase CmAP (0.3 mg/mL, 2300 U/mg towards *p*NPP), with the use of LPS as a substrate, was determined by quantitative analysis of phosphorus remaining in LPS after hydrolysis. An aliquot of the enzyme (0.0003 – 0.04 mg/mL) in 1 M DEA, pH 10.3, and LPS samples were mixed in glass vials with caps (total volume 1 mL) and incubated at 37 °C with stirring for 30 min, 2 h, and 12 h. After the end of reaction, the samples were transferred into dialysis tubes (with a pore size of 3000 Da) and dialyzed against distilled water for 2 days in a cold room for removal of free phosphorus.

For quantitative determination of phosphorus in the LPS solution before and after treatment with the alkaline phosphatase CmAP, a universal molybdate reagent was used [35], for which a working solution was prepared: 26 mL of 1 N sulfuric acid was added to 5.5 mL of the initial reagent and the volume of distilled acid was adjusted to water up to 100 mL. An aliquot of the LPS solution (200 μL) was taken into a test tube and evaporated to dryness in a heating oven at 100°C. Then, 0.05 mL of 72% perchloric acid was added to the dry residue and burned in a duralumin block at 180-200 °C for 20 min. After cooling, 0.45 mL of the working reagent was added to the test tubes. The mixture in the test tube was thoroughly mixed using a vortex, and the test tubes were placed in a boiling water bath for 15 min. After the formation of phosphomolybdenum blue, the tubes were cooled and the optical density of the samples was measured in a quartz cuvette (l = 1 cm) at 815 nm on a Specol spectrophotometer (Carl Zeiss, Germany). For each sample, 3 parallel measurements were made. For each measurement, a control sample was used (buffer without LPS with the same concentration of the enzyme CmAP), the absorbance of which did not exceed 0.04-0.05 optical density units.

A calibration curve for determining phosphorus in the LPS samples was drawn using monosodium phosphate (NaH_2_PO_4_ x 2H_2_O) (AppliChem, Germany) dissolved in water (stock solution - 7.8 mg in 250 mL) in the concentration range 0.03 - 0.3 μg /mL.

### 2.4. Phylogenetic and biosynthetic gene clusters analyses

For identifying the homologues of the *C. amphilecti* KMM 296 alkaline phosphatase CmAP (accession no. KGA01942), global blast, blastn, and tblastn (Available online: https://blast.ncbi.nlm.nih.gov/Blast.cgi, accessed on 18 August 2023) were performed.

For the ALP phylogeny, 36 publicly available *Cobetia* spp. genomes were used to search for the genes (CDS) encoding for the homologues of the *C. amphilecti* KMM 296 alkaline phosphatase CmAP (accession no. KGA01942), as well as other genes and gene products related to the functional annotation “alkaline phosphatase” (Table S1, sheet “ALP types”). Using own R scripts for processing tabular data (libraries: rtackleyer, readr, plyr, dostats, rtracklayer, sybil, data.table), the *gtf* of all *Cobetia* strains were analyzed to create a list of genes described as “alkaline phosphatase” (Table S1, sheet “ALP types”). In the *gtf* files there are 3 kinds of description of alkaline phosphatase: “alkaline phosphatase”, “alkaline phosphatase D family protein”, “alkaline phosphatase family protein”. The found protein IDs were used for multiple alignment and phylogenetic tree construction using Clustal and ITOL (Available online: https://itol.embl.de/; accessed on 24 October 2023), respectively. In addition, the referent ALP proteins of the family PhoA were taken in the analysis, namely: *Homo sapience* (NP_001622.2, P09923.2), *Euprymna scolopes* (AER46070, AER46069), *Moritella* sp. 5 (QUM82918), *Vibrio* sp. G15-21 (Q93P54), and *E. coli* (NP_414917, NP_311634, NP_417233). The referent ALP proteins of the family PafA were of *Flavobacterium* spp.: WP 073408805; WP 011921542, WP 149206384, WP 198858267, WP 091133708. The gene products with the annotation “alkaline phosphatase D family protein” were taken as the family PhoD ALPs (Table S1, Table S2). The unclassified proteins with the functional annotation “alkaline phosphatase family protein” were referred to the family PhoX or PafA, which were manually checked and compared with literature data (Table S1, Table S2). Using gene_neighbor.R and Integrative Genomic Viewer (IGV) browser (Available online: [IGV](https://igv.org/app/); accessed on 23 July 2023), the metG gene neighboring clusters containing the CmAP homologues (KGA01942) were identified in all gtf for the analysis of their location in each strain chromosome studied (Table S1, sheet “ALP neighbors”).

The evolutionary history was inferred by using the Maximum Likelihood method and Jones et al. w/freq. model [36]. The tree with the highest log likelihood (−22741.47) was taken. The initial trees for the heuristic search were obtained automatically by applying Neighbor-Join and BioNJ algorithms to a matrix of pairwise distances estimated using the JTT model, and then selecting the topology with superior log likelihood value. The analysis involved 137 amino acid sequences. There was a total of 880 positions in the final dataset. Evolutionary analyses were conducted in MEGA 11 [37].

To search for metabolic pathways involving the homologues of the *C. amphilecti* KMM 296 alkaline phosphatase CmAP (accession no. KGA01942), the program gapseq (https://github.com/jotech/gapseq, accessed on 23 July 2023) was used to build genome-wide metabolic network models using a modified automatically updated ModelSEED biochemistry database [38]. Prediction of pathways, transporters and complexes is based on a protein sequence database derived from UniProt and TCDB.

To visualize the biosynthetic gene cluster synteny, clinker, a Python-based tool, and clustermap.js, a companion JavaScript visualization library, were applied that together allowed automatically generating accurate interactive gene cluster comparison figures directly fro m sequence files [39]. Clusters were identified using antiSMASH (https://antismash.secondarymetabolites.org/, accessed on 09 December 2023), then the resulting .gbk sequences were aligned using clinker. The cluster alignments created using clinker are visualized using clustermap.js and edited using Adobe Illustrator [40].

## 3. Results and Discussion

### 3.1. Dephosphorylation activity of the recombinant alkaline phosphatase CmAP towards E. coli LPS

The *E. coli* LPS were used as the P-containing substrates for the *C. amphilecti* KMM 296 alkaline phosphatase CmAP (accession no. KGA01942), with a high ALP specific activity and catalytic efficiency relatively to other bacterial homologues, in order to study its physiological function and substrate specificity in nature [3,10,15,33,28].

The bacterial LPS recognition and dephosphorylation is a very ancient function of ALPs and belongs to the innate immunity system of multicellular organisms [10]. Alterations in the expression and activity of LPS-dephosphorylating ALP of the light organ (photophore) of the squid *Euprymna scolopes* regulate colonization and abundance of its luminescent symbiont *Vibrio fischeri* [11]. The increase in the level of dephosphorylated LPS by sunset is a signal for the bacteria to increase population size and consequently photophore luminescence intensity, and protect simultaneously the invertebrate from excessive inflammation and destruction of its own tissues by the endotoxic phosphorylated lipid A [11]. The possible effect of the bacterial ALP on LPS has not been investigated except for the finding that exogenous *E. coli* ALP, which was applied to mice infected with *Pseudomonas aeruginosa*, positively modulated the growth of commensal bacteria in the animals [41]. This probably led to increased competitiveness of their microflora with the infectious agent and reduced its production of enterotoxins [41].

LPS is the main component of the outer membrane of Gram-negative bacteria, occupying about 70% of its surface [42]. The complete structure of the amphiphilic LPS molecule, the smooth type of LPS (S-LPS), characteristic of most wild-type bacterial strains found in nature, consists of three parts: the hydrophobic lipid A, core oligosaccharide, and O-polysaccharide chain. The O-polysaccharide is attached to the core oligosaccharide, which in turn is bound to the lipid A moiety. Lipid A of most of the studied bacteria is a dephosphorylated *β*-1,6-linked glucosamine disaccharide (diaminogenciobiose) bearing higher fatty acid residues. In addition to the S-form of LPS, bacteria also synthesize core-defective LPS molecules (R-chemotypes), which have lipid A and a core part of different lengths. The core-defective structures are labelled as Rb - Re-chemotypes, and the structures with a complete outer core as Ra-chemotypes [43,44]. In the LPS micelles formed in aqueous solutions, lipid A is located inside the micelles, and the O-polysaccharide, constructed in most cases from hydrophilic monosaccharide residues as in bacterial cells, is directed towards the aqueous phase. Therefore, the solubility of LPS depends on the length of O-polysaccharide [42,44].

Thus, the amount of detectable phosphorus in the LPS sample solubilized in 0.1 M Tris-HCl buffer, pH 7.7, increased 6-fold after reducing the LPS concentration from 1 to 0.2 mg/mL and increasing the incubation temperature to 37 °C (data not shown). It was significantly increased after the addition of TEA and, consequently, alkalization incubation medium to pH 10.0 (sample N 3) compared to the LPS sample N 1, (LPS incubation at 24 °C and without an addition TEA). The maximum phosphorus content was determined in the LPS sample N 3 (Figure 1).

**Figure 1.**
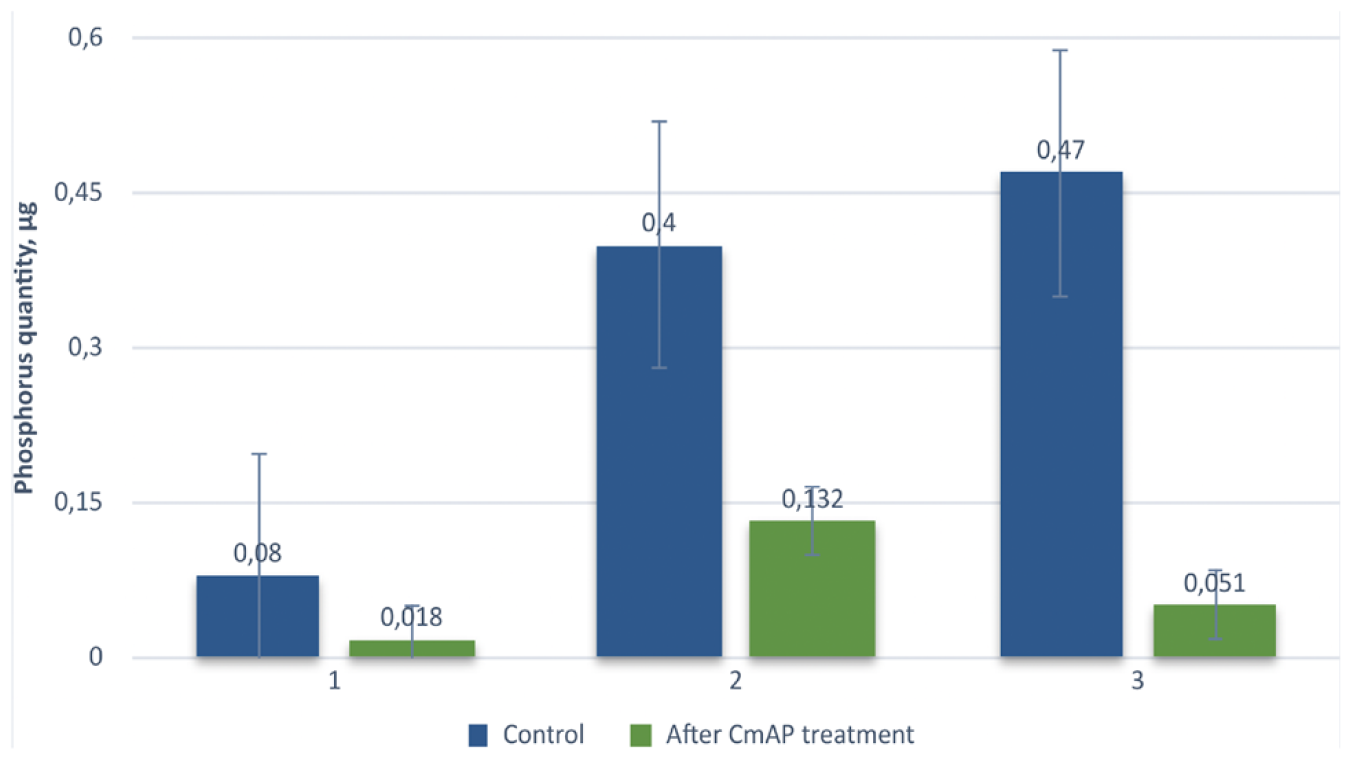
The phosphorus quantity (μg) in the *E. coli* 055:B5 LPS (samples N 1-3) before (control, blue) and after (green) treatment with 0.024 mg/mL recombinant alkaline phosphatase CmAP (2300 U/mg) for 30 min at 37 °C with the use of 0.2 mg/mL LPS samples dissolved in 200 μL of: **1** - 0.1 M Tris-HCl buffer, pH 7.7, at 24 °C for 12 h; **2** - 0.1 M Tris-HCl buffer, pH 7.7, at 37 °C for 12 h; **3** - 0.1 M Tris-HCl buffer, pH 7.7, with an addition of triethylamine (TEA) at 24 °C for 12 h.

Expectedly, the maximum dephosphorylation activity of the recombinant enzyme CmAP was observed against the LPS sample N 3, for which the amount of hydrolyzed phosphorus was 89% of the control sample (LPS non-treated with the alkaline phosphatase CmAP) (Figure 1). The reaction time at the optimal conditions should be at least 30 min at the protein concentration 0.024 mg/mL of the CmAP preparation with the specific pNPP-dephosphorylation activity 2300 U/mg protein (Figure 1).

Thus, the alkaline phosphatase CmAP of the marine bacterium the *C. amphilecti* KMM 296 exhibits the enzymatic activity toward the *E.coli* LPS, which depends largely on the degree of their availability of phosphate residues in the LPS molecule in a solution (disaggregation state). The enzyme CmAP is able to dephosphorylate LPS almost completely, if it has the complete structure (S-form) and is pre-incubated for 12 h at 37 °C (Figure 1).

The specificity of the marine bacterial ALP CmAP towards LPS is similar to the epithelial enzymes of eukaryotes, in particular the human isoenzyme IAP, which dephosphorylates LPS of its own and pathogenic intestinal microflora to prevent inflammation and persistence of bacteria and endotoxins into the bloodstream [4,6-8,10].

The findings on the LPS-dephosphorylation activity of the marine bacterial enzyme CmAP provide a promising basis for the development of a novel therapeutic approach to neutralize the effects of bacterial endotoxins, such as Crohn’s disease, sepsis, and endotoxic shock.

### 3.2. Multiplicity of alkaline phosphatase phylotypes in the species Cobetia amphilecti

#### 3.2.1. Structural classification of the bacterial alkaline phosphatases

Bacterial ALPs can be divided by the amino acid sequence (primary structure) into three main structural families: PhoA, PhoD and PhoX belonging to COG1785, COG3540 and COG3211 proteins, respectively, which originate from different ancestral genes according to the classification of the Clusters of Orthologous Genes (COG) database [26]. The structure of the PhoA family enzymes was the first to be studied, as the classical phosphomonoesterase of *E. coli* belongs to it [10]. Subsequently, other enzymes non-homologous to the *E. coli* alkaline phosphatase, but with similar physiological functions, were discovered in the environmental bacteria [13,14,22,23,31]. The high variability of protein sequences in the alkaliphilic enzymes is characteristic both for representatives of different families and within each family in dependence on the taxonomic affiliation of their microbial producers [23,26]. Despite the low sequence identity among representatives of the PhoA, PhoD, and PhoX families, their main common property is the production of P_i_ during the depletion of its stores in the environment, and, consequently, their expression and enzymatic activity are inhibited by the high P_i_ concentrations according to a feedback reply [12,14]. However, unlike the Mg^2+^-activated and Zn^2+^-containing phosphomonoesterases of the PhoA family, the members of the PhoD and PhoX families exhibit their maximal activity in the presence of Ca^2+^/Co^2+^ ions against the wider range of substrates, both phosphomono- and diesters [19,23,31]. In addition, some alkaline phosphatases/phosphodiesterases PhoD contain Fe^3+^ in their bimetallic active centers instead of Zn^2+^ [31,45]. Such an unusual architecture of the active centre and its Fe^3+^-specificity could have evolved in the zinc-deficient as an adaptation of the bacterium to survive in a particular environment. This property may have been acquired by soil bacteria from a plant purple acidic phosphatase PAP [45].

Some authors include in the classification of bacterial ALPs a fourth family of constitutive highly active Ca^2+^-dependent enzyme PafA, with a broad substrate specificity and unknown metabolic function, common in the genomes of flavobacteria, mainly *Bacteroidetes*, associated with plant rhizospheres, because their expression and enzymatic activity are not inhibited by the reaction product, P_i_, and are not controlled by known regulators unlike other ALPs [14,26]. The presence of non-inducible and unrepressed PafA alkaline phosphatase in flavobacteria has been shown to promote rapid remineralization of various organophosphates and P_i_ accumulation, which enables a secondary growth of other bacterial species in microbiomes [14]. In addition, the PafA-like ALPs can be active against phosphodiesters, expanding the role of these enzymes in nature [14,18]. However, some representatives of the PhoD, PhoX, and PhoA families may exhibit the similar properties [13,14,18,19]. The highly active and catalytically efficient enzyme CmAP isolated from the marine bacterium *C. amphilecti* KMM 296 was also not inhibited by a large concentration of P_i_ [15,33]. A recombinant PhoA analogue from the marine bacterium *Alteromonas mediterranea* showed a broad substrate specificity towards phosphodi-, phosphotriesters, and sulfates at low substrate concentrations, whereas the enzyme exhibited a high phosphomonoesterase catalytic efficiency at the high substrate concentrations [13]. Thus, the structural classification of the ALP enzymes only reflects their belonging to different homologous lineages, descended from different ancestral genes that evolved independently to perform the same functions in the organism and/or microbiome [22,23,26].

#### 3.2.2. Searching for homologues of the *C. amphilecti* KMM 296 alkaline phosphatase CmAP

According to the blast-based searching against the sequence query KGA01942 encoding for the *C. amphilecti* KMM 296 alkaline phosphatase CmAP, it is homologue for the PhoA *E. coli* ALP. The multiple alignment showed that the PhoA sequences of *E. coli* are differ significantly from the PhoA of *C. amphilecti*, with overall identity 32.21% at the cover 61% (Figure S1). The search for the KGA01942 protein among the *Cobetia* spp. genomes (Table S1, Table S2) identified the eigth homologous sequences in eight from 35 strains, indicating that it is characteristic for the strains of the subspecies *C. amphilecti* (Table S1, Table S2) according to the gene-specific and genome-based classification of the *Cobetia* genus as described earlier [31,46]. Although both groups of the strains, *C. amphilecti* and *C. litoralis*, belong to the same phylotype in terms of the whole genome characteristics [46], the species is divided into two subspecies based on the content of alkaline phosphatases (Table 1, Table S1). Probably, this divergence emerged when the parent population had changed the lifestyle from free-living to host-associated [31].

**Table 1.** The content and distribution of alkaline phosphatase families in the *Cobetia* spp. isolates

| Original Strain title | Actual Species* | Accession ID | ID protein | ALP family | Isolation source |
| --- | --- | --- | --- | --- | --- |
| *C. marina* JCM 21022<sup>T</sup> | *C. marina* | GCF_001720485.1 | WP_240499655.1<br>WP_084208519.1<br>WP_240499593.1 | PhoD<br>PhoX<br>PhoD | Littoral water sample (USA: Woods Hole, MA) |
| *C. pacifica* NRIC 0813<sup>T</sup> | *C. marina* | GCA_030010515.1 | WP_284728477.1<br>WP_240704267.1<br>WP_084208519.1 | PhoD<br>PhoX<br>PhoX | Sandy sediment (Russia: The Sea of Japan) |
| *C. litoralis* NRIC 0814<sup>T</sup> | *C. amphilecti* | GCF_029846315.1 | WP_249330383.1<br>WP_279830791.1<br>WP_279833222.1<br>WP_279832006.1 | PhoD<br>PafA<br>PhoX<br>PhoD | Sandy sediment (Russia: The Sea of Japan) |
| *C. amphilecti* NRIC 0815<sup>T**</sup> | *C. amphilecti* | GCA_030010415.1 | WP_284726718.1<br>WP_284726808.1<br>WP_284726995.1<br>WP_284727213.1<br>WP_284727576.1 | PhoD<br>PhoA<br>PhoX<br>PhoD<br>PafA | The finger sponge *A. digitatus* (The Sea of Okhotsk, Sakhalin Island, Piltun Bay) |
| *C. amphilecti* KMM 296<sup>**</sup> | *C. amphilecti* | GCF_000754225.1 | WP_043332298.1<br>WP_043333989.1<br>WP_245163010.1<br>WP_043336117.1<br>WP_052384691.1 | PafA<br>PhoD<br>PhoA<br>PhoX<br>PhoD | Coelomic fluid of mollusc *C. grayanus* (Russia: The Sea of Japan) |
| *C. crustatorum* JO1<sup>T**</sup> | *C. crustatorum* | GCF_000591415.1 | WP_248623642.1<br>WP_282705494.1<br>WP_282705495.1<br>WP_024952594.1 | PhoA<br>PhoD<br>PhoD<br>PhoX | Fermented shrimp (South Korea: Daejeon) |

\*According to Nedashovskaya et al., 2023; \*\* The strains containing the PhoA family.

Remarkably, the mollusc strain *C. amphilecti* KMM 296 and the coral-associated type strain *C. amphilecti* NRIC 0815^T^, as well as other available *C. amphilecti-*like isolates, have been found to possess 5 proteins with the functional annotation “alkaline phosphatase”, including the phoA family ALP, in contrast to 4 ALPs found for the *C. litoralis*-like strains (Table 1, Table S1, Table S2). In addition, the PhoA gene was found in the type strain *C. crustatorum* JO1^T^ from a shrimp and two strains *C. crustatorum* SM1923 and *Cobetia* sp. QF-1 (Table S2), which were suggested to belong to the new species of the genus *Cobetia* [46]. However, the species *C. crustatorum* and two new species are phylogenetically distant from other *Cobetia* species, therefore they are easily classified using 16S RNA gene analysis, unlike the closely related and undistinguished between each other isolates of the *C, amphilecti* and *C. litoralis* or *C. marina* and *C. pacifica*, respectively [31,46].

The nucleotide substitutions in the PhoA family coding sequences were detected in the *C. amphilecti*-like genomes (∼1%) and in the genomes of *C. crustatorum* JO1^T^ (∼11%) and *Cobetia* sp. SM1923 (∼11%) (Table 2). The substitutions do not hit the active sites of the enzymes except for *C. crustatorum* JO1^T^, which has down start (+45 amino acids) the missing key position 21D (a catalytic Asp), indicating a non-functional homologue of the enzyme CmAP [33]. However, the exact metabolic function of the enzyme CmAP (KGA01942) and its homologues was not identified by the program gapsec except for the pathway producing NADH from the consumed NADPH, mostly associated with the intracellular 5’-nucleotidase (surE), probably, due the low identity of the model enzymes, which participation in metabolic reactions is verified [38]. Gapseq detected pathways associated with ALP in all strains because of the blast-based homologs of the desired enzyme are found in the genomes of all bacteria studied, but they are short (<100 amino acids) and with low homology (Expect > 1, Identities <50%).

**Table 2.**
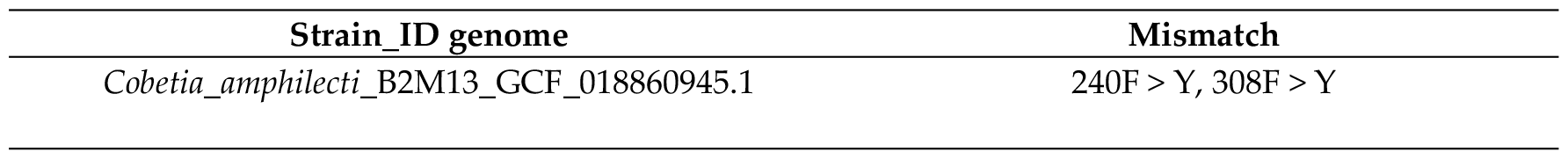
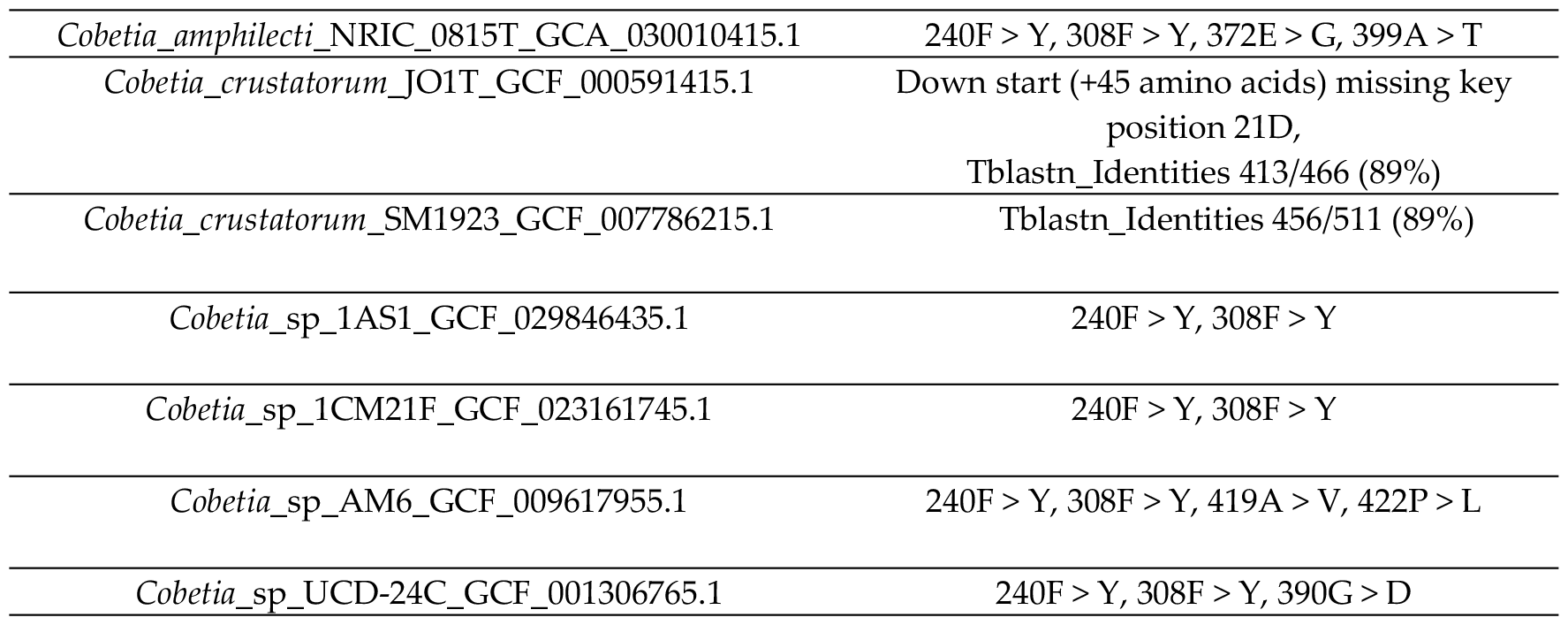
Comparative analysis of the PhoA family structures found in the *Cobetia* genomes

Using the IGV browser, an unnamed gene with an alkaline phosphatase product homologous to the *C. amphilecti* KMM 296 CmAP (KGA01942) was identified in gtf files of ten *Cobetia* strains, which predominantly located next to the metG gene (+/- 2 positions). In this regard, we decided to isolate the region centered on the metG gene. The Table S1 (sheet “ALP neighbours”) was created based on the results of the analysis. Each cell contains a gene and a product in the format “gene|product” (e.g.: “metG |methionine-tRNA ligase”). The CmAP-like gene (KGA01942) is present in the table as “NA |alkaline phosphatase” because the gene name is not labelled in the map. However, the non-homologous ALP of the *C. pacifica* NRIC 0813^T^ is also located in the similar metG-cluster (Table S1). In addition to the methionine-tRNA ligase (metG), in the immediate environment there are electron transport complex subunits RsxD, RsxC, RsxB, RsxA, DUF3465 domain-containing protein (Si-specific NAD(P)(+) transhydrogenase), TerB family tellurite resistance protein, unnamed alkaline phosphatase, cell wall hydrolase, RNA pseudouridine synthase, double apbC genes and iron-sulfur cluster carrier protein ApbC playing a role in regulating NADH oxidation, dCTP deaminase, autotransporter domain-containing SGNH/GDSL hydrolase family protein (proteases and lipases related) (Table S1, sheet “ALP neighbors”). Such gene clusters are demand in electron transfer reactions, respiration, DNA repair, gene regulation and, consequently, may relate to a rearrangement of the cell wall and cellular phenotypes under oxidative stress or iron/sulfur limitation in both Gram-negative and Gram-positive bacteria [47, 48]. Besides P-nutrient scavenging under P_i_ deficiency, PhoA also functions, under the P-replete condition, to constrain secondary metabolites biosynthesis, and cell division. These functions have important implications in maintaining metabolic homeostasis and preventing premature cell division [49]. These facts might explain the biofilm-degrading behavior of many different bacteria after their treatment by the PhoA alkaline phosphatase CmAP solution [28]. Although only 10 strains have the CmAP-like gene in the immediate neighborhood of the metG (Table S1, sheet “ALP neighbors”), the similar extended metG-regions for the different strains are observed. It should be noted that in the genomes of *Cobetia* sp. UCD-24C (GCF_001306765.1) and *C. crustatorum* JO1^T^ (GCF_000591415.1), the CmAP-like genes (like KGA01942) locate not in the neighborhood of the metG gene (Table S1). In the genomes of *Cobetia* sp. UCD-24C (GCF_001306765.1), *C. crustatorum* JO1^T^ (GCF_000591415.1), *C. marina* NBRC 15607 (GCF_006540105.1), and *Cobetia* sp. 10Alg146 (GCF_029846385.1), the metG environment is significantly different.

#### 3.2.3. Phylogenetic analysis of the alkaline phosphatases in the genus *Cobetia*

The phylogenetic analysis of the found *Cobetia* spp. ALPs revealed that each *Cobetia* strain contains from 2 to five ALPs belonging to the different structural families, of which two different PhoD or two different PhoX structures are necessarily present (Table 1, Figure 2, Table S2, Figure S2).

**Figure 2.**
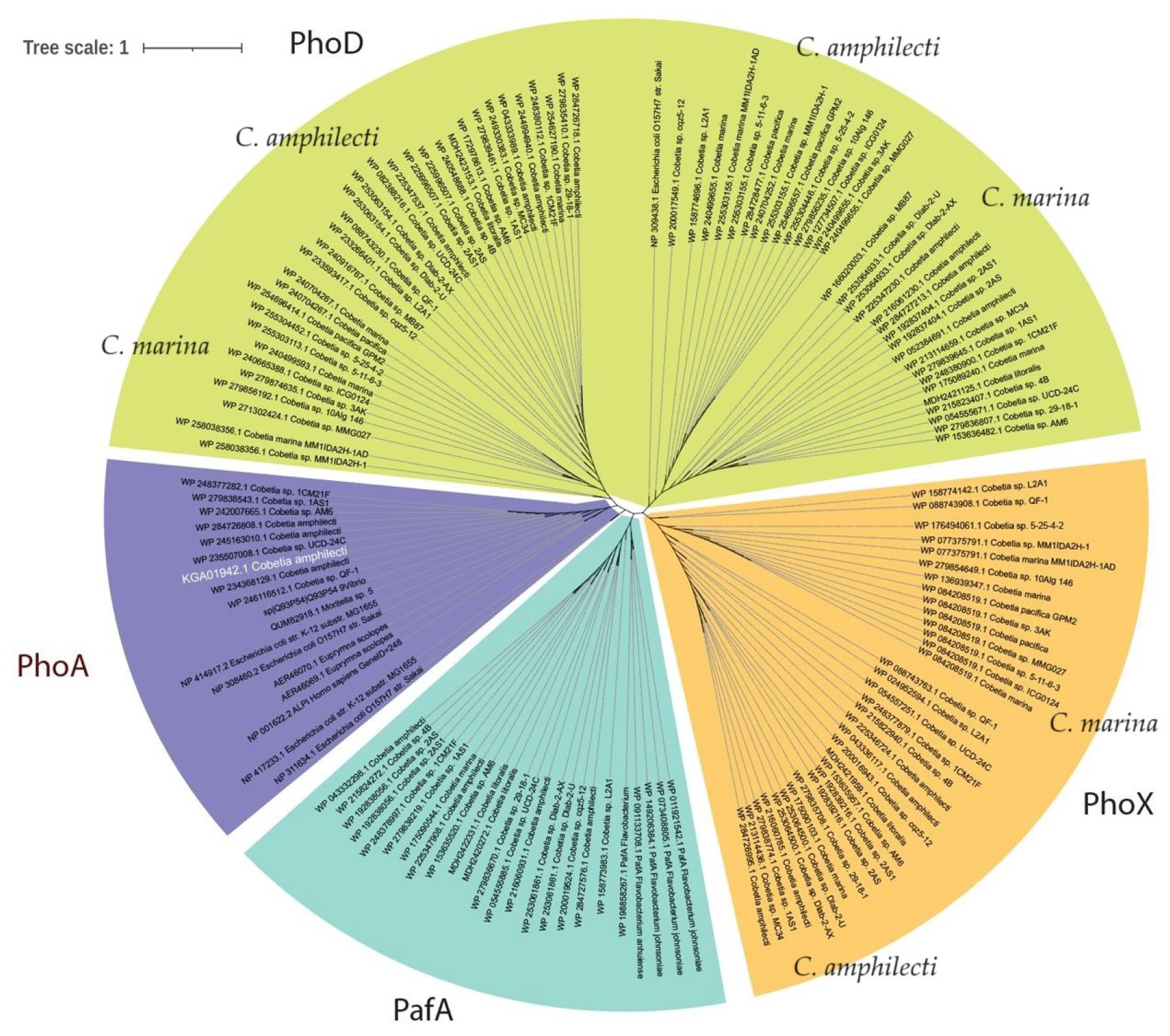
Phylogenetic tree based on the similarity of full-length amino acid sequences of the *Cobetia* spp. alkaline phosphatases constructed using the MEGA 11 program by the maximum likelihood method. GenBank database access numbers are indicated before the species titles of the strains. Bootstrap values at the nodes of the bootstrap consensus tree constructed from 1500 samples are at least 50%. Alkaline phosphatase structures are grouped into the protein families: 0, 1 - alkaline phosphatase PhoD; 2 - alkaline phosphatase PhoA; 3 - alkaline phosphatase PafA; 4 - alkaline phosphatase PhoX. The *C. amphilecti* KMM 296 alkaline phosphatase CmAP (accession no. KGA01942). The bootstrap values at the nodes of bootstrap consensus tree based on 1,500 samplings are not less than 50%. The AP structures are clustered into the protein families: alkaline phosphatase PhoD (green); alkaline phosphatase PhoX (orange); alkaline phosphatase PafA (turquoise); alkaline phosphatase PhoA (purple): CmAP (white).

Each PhoD or PhoX structure from one strain seems to belong to the different lineage but to be species-specific according to their clusterization into the species-based groups within one homologous lineage (Figure 2). For example, the strain *C. amphilecti* KMM 296 has two PhoD sequences under the accession numbers WP_043333989.1 and WP_052384691.1 (Table 1), which located on the left and right PhoD branches of the phylogenetic tree, indicating their origination from the different parent genes (Figure 2). The *C. marina* strains also possess two PhoD genes located in the different PhoD branches, each of which differs from the sequences of *C. amphilecti* of the same homologous lineage. Moreover, the addition of the referent sequences from *Flavobacterium* spp. belonging to the novel ALP family PafA [14,26] allowed identifying the Paf-like proteins in the *Cobetia* isolates, which were clusterized in the separate branch (Figure 2, Figure S2) consisting of other ALP enzymes that coincide with the unclassified alkaline phosphatases from the Table S1 (under the number 2, colored in red, sheet). Remarkably, the PafA family was found only in the genome of the species *C. amphilecti*, including both subspecies, *C. amphilecti* and *C. litoralis* (Table 1, Table S2). This indicates that the *C. amphilecti* strains may colonize plant roots similarly to the PafA-containing flavobacteria [14]. Indeed, the strain *Cobetia* sp. UCD-24C was isolated from the *Zostera* spp. roots and was found to possess the PhoA family gene as other *C. amphilecti*-like isolates (Table S2).

It is evident that the PhoA sequences of the *Cobetia* strains are located in the cluster common for the LPS-detoxifying ALPs of human and squid (Figure 2, Figure S2). The PhoA structural family of ALPs is predominantly associated with marine heterotrophic bacteria of *Bacteroidetes* and γ-*Proteobacteria* neighboring cyanobacteria populations [13,22], as well as the microorganisms isolated from plant and animal microbiomes, such as a symbiont (pathogen) *Vibrio* spp. of planktonic crustaceans (Copepods), a symbiont (facultative pathogen) of human *E. coli*, a sugarcane root endophyte *Enterobacter roggenkampii* [29,50,51], marine fish pathogens *V. splendidus* [52], and *Moritella* sp. [53]. In addition, most eukaryotes have their own PhoA enzymes, with the exception of some plants, to modulate their microbiomes and tissue inflammation [9-11]. Probably, the pathogenic or symbiotic bacteria have evolved the strategy to evade host immunity, including the acquisition of the immunomodulatory PhoA alkaline phosphatase [53]. It follows that all PhoA alkaline phosphatases belong to a common homologous protein lineage [10] and are clustered into a separate branch of the phylogenetic tree. In contrast to the species *C. amphilecti* (Table 1), the ALP content of the subspecies *C. marina* and *C. pacifica* is restricted only by three genes of the families PhoD and PhoX (Table S1, Table S2).

Alkaline phosphatases of both families are widely distributed in marine and soil bacteria. However, the phoD family is more adapted to the lifestyle of soil bacteria, whereas the phoX family is more characteristic of marine bacteria, including cyanobacteria, and in the soil microbial community in response to an increase in total organic matter content. In this regard, the phoX genes dominate in the marine metagenomic databases [22,23]. Whereas, the presence of the PhoA family structures in the marine metagenomes may be an indicator of a large number of *Bacteroidetes* representatives, which were abundant during phytoplankton blooms [13,22]. In addition, most bacterial genomes are characterized by a single ALP family in the genome, whereas a single bacterium rarely has both members of the PhoX and PhoA or PhoX/PhoA and PhoD families [22]. However, thirty-five per cents of the owners of multiple paralogues of ALPs have at least one gene of the PhoA family, whereas 22 and 17% of the bacterial genomes have PhoX and PhoD genes, respectively [24]. The complete range of biological functions of paralogues encoding for alkaline phosphatase isoenzymes in a single organism remains still unknown. The differences in structural and, consequently, physicochemical properties of the ALP paralogues suggest their functional diversification during assimilation of different organophosphates under conditions of phosphorus deficiency and different chemical parameters of the environment [13,24].

### 3.3. Biosynthetic gene clusters analysis in the genus Cobetia

The analysis of biosynthetic gene clusters of the type strains of the genus *Cobetia* confirmed the species demarcation. Considering the main biosynthetic gene clusters (BGCs) inherent in the genus *Cobetia*, a high level of similarity of both the arrangement of genes in clusters and the sequences of individual genes was found. Despite a high BGC similarity between the all *Cobetia* isolates, belonging *C. litoralis* and *C. pacifica* to the species *C. amphilecti* and *C. marina* respectively [46] is also evident at the level of genus-specific BGCs. In particularly, the ectoine BGCs allow dividing these the species *C. amphilecti* and *C. marina* by decreasing sequence identity of the marker sensor genes encoding for diguanylate cyclase and phosphodiesterase (Figure 3). This signaling pathway based on the bacterial sensitivity to the concentration of cyclic dimeric GMP (c-di-GMP), a universal second messenger, in order to control the cell growth and secondary metabolites production such as, apparently, the biosynthesis of ectoine in a high-salt medium [31,54].

**Figure 3.**
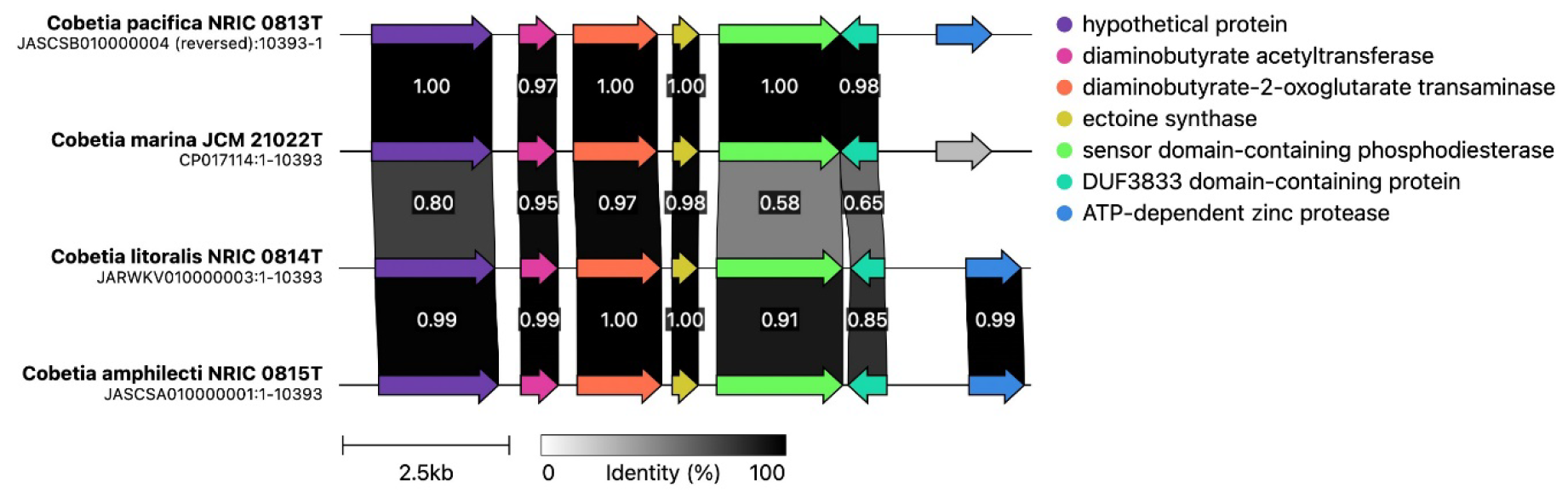
The ectoine BGCs in the type strains of the genus *Cobetia*. The different identity between the genes is reflected by gradient of gray, the identical genes are in black.

The genus-specific BGC encoding for the Ni-siderophore synthesis genes was found to be indistinguishable between the *Cobetia* isolates (Figure S2), indicating the high significance of the iron uptake, when bacteria grow under oxidative stress or iron limitation, particularly, for inclusion as cofactor in the active sites of the specific enzymes, such as alkaline phosphatase/phosphodiesterase PhoD in *C. amphilecti* KMM 296 [31,45,47]. However, the genes of aspartate aminotransferase, monooxygenase, and sigma 70 family PNA polymerase factor were absent in the *C. amphilecti* KMM 296 cluster (Figure S2). Contrarily, the T1PKS BGC seems to be highly species and strain specific (Figure 4).

**Figure 4.**
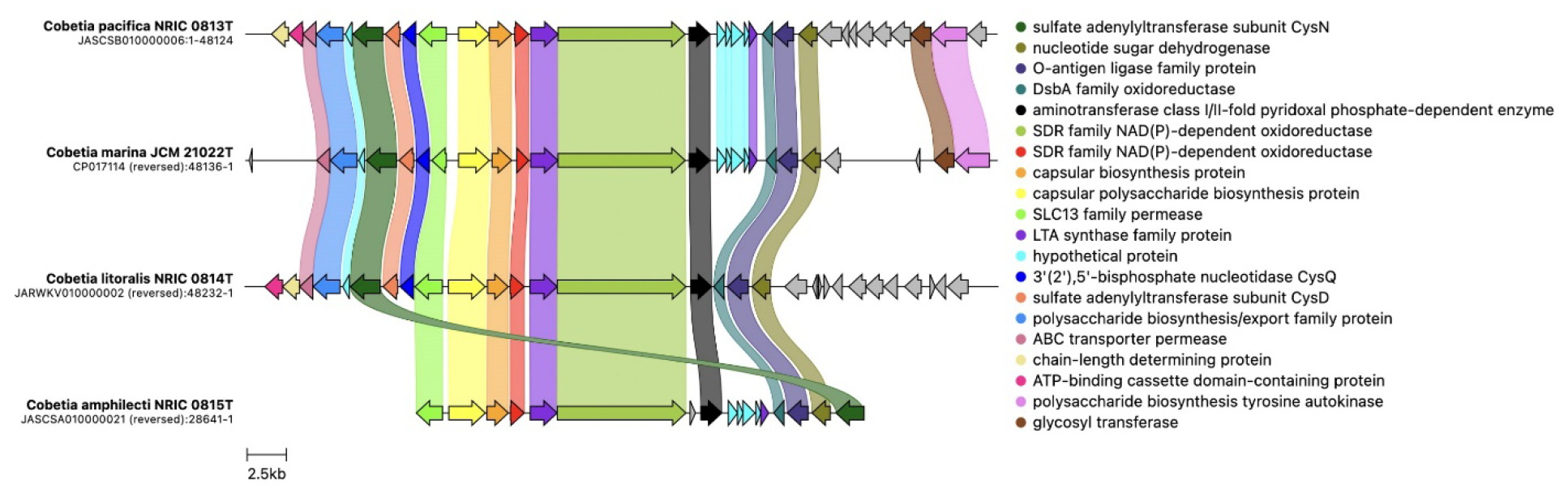
The T1PKS biosynthetic gene clusters in the type strains of the genus *Cobetia*.

However, several key genes (capsular polysaccharide biosynthesis protein, SDR family NAD(P)-depended oxidoreductase, type I polyketide synthase, DsbA family oxidoreductase) encoding for the synthesis of extracellular polysaccharides of the high conservativity are found in all genomes and their divergence also coincide with the genome-based taxonomic classification for the genus *Cobetia* [46]. It has been determined that the strains *C. pacifica* KMM 3789^T^, KMM 3878, and *C. litoralis* KMM 3880^T^ produce sulfated O-polysaccharides (OPS) composed of trisaccharide repeating units and include D-glucose 3-sulfate and D-galactose 3-sulfate, D-galactose 2,3-disulfate, and 2-keto-3-deoxy-D-manno-octanoic acid 5-sulfate, respectively [55-56]. The *C. pacifica* capsular polysaccharides were found to exhibit an antiproliferative effect and suppress the colony formation of DLD-1 and MCF-7 cells in a different manner [55]. Whereas, the LPS and O-deacetylated OPS from *C. litoralis* KMM 3880^T^ inhibited a colony formation of human melanoma SK-MEL-28 and colorectal carcinoma HTC-116 cells [56]. In this regard, it is of interest to study the polysaccharide structures of all *Cobetia* strains, particularly the *C. amphilecti* LPS to determine the length and content of acyl chains present on lipid A. The ability of the PhoA enzyme CmAP to cleave own LPS may shed light on its role for the marine bacterium survival, because the strains of *Moritella* spp. containing lipid A with the highest amount of C16 acyl chains and the PhoA family ALP is found to be immuno-silent for their host [53].

## 5. Conclusions

The alkaline phosphatase of *C. amphilecti* KMM 296 belong to the PhoA structural family similarly to eukaryotic enzymes, for which the LPS-dephosphorylating and detoxifying function were confirmed. Probably, some bacteria like the immuno-silent *Moritella* spp. may use the PhoA enzymes for imitation of the host ALP to manage other competing bacteria of the host microbiome and to hide from the immune system.

However, this hypothesis is still to be verified. Nevertheless, the abundance of different families ALPs in the *Cobetia* isolates indicates their ability to survive in different ecological niches with P-deplete or replete medium because of capability of degrading many types of organophosphates. This may provide other members of an ecological niche including plants such essential macronutrient as phosphorus. The ability to produce ectoine, a variety of sulfated exopolysaccharides, and the PhoA type alkaline phosphatases with LPS-dephosphorylating activity provide the *Cobetia* species survival in the high salty and competing conditions, as well as a promise of their use in biotechnology and medicine.

## Supplementary Materials

The following supporting information can be downloaded at: www.mdpi.com/xxx/s1, Figure S1: Multiple sequence alignment for the PhoA family alkaline phosphatases of *C. amphilecti* KMM 296 CmAP and *E. coli*; Figure S2: The iTOL phylogenetic tree of the *Cobetia* spp. genes encoding for alkaline phosphatases based on the whole-genome computational analysis; Figure S3: Ni-siderophore biosynthetic gene clusters for the species *Cobetia amphilecti*; and Table S1: Whole-genome based analysis of alkaline phosphatase genes and their localization in *Cobetia* spp.; Table S2: The content and distribution of alkaline phosphatase families in *Cobetia* spp.

## Author Contributions

Conceptualization, L.A.; methodology, L.B, S.B. and O.K.; validation, L.B. and S.B.; investigation, N.B., V.V. and Y.N.; resources, L.T.; writing—review and editing, L.B.; visualization, O.K. and G.B.; project administration, O.S.; funding acquisition, L.T. All authors have read and agreed to the published version of the manuscript.

## Data Availability Statement

Data available in a publicly accessible repository

